# Arousal-Driven Serial Dependence: The role of Internal States in Temporal Judgments

**DOI:** 10.1101/2025.06.15.659791

**Authors:** Si Cheng, Wenjie Jin, Xuanbing Shan, Elif Bengisu Kara, Sabiha Ecenur Bilgiç, Svetlana Salomasova, Quirin Unterguggenberger, Zhuanghua Shi

**Affiliations:** General and Experimental Psychology, Department of Psychology, LMU Munich, Munich, Germany; Chair of Neuro-cognitive psychology, Department of Psychology, LMU, Munich, Germany

**Keywords:** Serial dependence, Time Perception, Arousal, Internal state

## Abstract

Emotional experiences shape not only how we perceive the present but also how recent experiences inform our current judgments. This study examined whether emotional arousal and valence modulate serial dependence in time perception—the tendency for prior stimuli to bias ongoing estimates. Participants viewed affective images that induced high-arousal (positive or negative) or low-arousal (neutral) states and then reproduced each image’s duration. High arousal reliably lengthened perceived durations. More critically, it strengthened serial dependence, with the strongest biases emerging when high-arousal trials followed other high-arousal trials, showing a state-dependent amplification of temporal integration. Emotional valence, by contrast, exerted no measurable influence on either temporal distortion or sequential bias. These findings show that emotional arousal tunes how the brain weighs past against present, highlighting internal emotional states as a key driver of temporal continuity in perception.

## Introduction

Serial dependence—our tendency to let recent experience bias current perception and decisions— has been documented across a wide range of visual judgements (for reviews, see Cicchini et al., 2023; Pascucci et al., 2023). Most work has focused on stimulus-driven factors such as uncertainty, contrast, or reliability (Cicchini et al., 2023), showing that observers lean more heavily on prior information when the present input is degraded or ambiguous (Fritsche et al., 2017; Pascucci et al., 2023). Much less is known about how internal emotional states modulate serial dependence.

Emotions bias interpretation, guide attention, and spill over into unrelated tasks (Schmidt & Schmidt, 2016). They are commonly described along two key dimensions: valence, which captures the pleasantness or unpleasantness of a feeling, and arousal, which reflects physiological activation (Bradley & Lang, 2017). Both dimensions distort the subjective flow of time. Positive and negative emotions alike lengthen perceived durations relative to neutral ones (Cui et al., 2023). Arousal, in particular, has been linked to a speeding of the internal clock, increasing the number of temporal “ticks” accumulated during an interval (Matthews & Meck, 2016).

Critically, time perception is also deeply warped by past experiences. Two robust biases illustrate this: central tendency, where perceived long durations shrink and short durations stretch toward the mean (see history of Vierordt’s law, Glasauer & Shi, 2021), and sequential dependence, where current estimates drift toward durations encountered moments before (Glasauer & Shi, 2022). Unlike visual processing, time perception lacks dedicated sensory receptors and relies more on internal cues - making it especially sensitive to fluctuations in attention, memory, and emotional states. Recent findings underscore this vulnerability. Increasing working memory loads within a trial–such as requiring observers to maintain multiple task-related features with a post-cue—amplifies sequential dependence (Cheng, Chen, & Shi, 2024). Similarly, Markov et al. (2024) showed that elevating load within the current trial strengthens serial dependence, whereas increasing load in the preceding trial weakened it. These results suggest that serial dependence emerges from a dynamic allocation of cognitive resources, which emotional states are well positioned to modulate.

Despite the extensive evidence that both emotion and recent experience bias perception, little is known about how they interact in terms of serial dependence. Emotional arousal influences attention, memory, and expectations (Phelps et al., 2014), all of which can recalibrate how prior information is weighted. If arousal alters quality or accessibility of present sensory information, the perceptual system may adjust its integration strategy, relying more or less on the immediate past to stabilize judgments.

The present study tested whether emotional arousal modulates temporal sequential biases in time perception. Participants viewed affective images from the International Affective Picture System (IAPS; Lang et al., 2018) that elicited high-arousal positive, high-arousal negative, or low-arousal neutral states. They then reproduced each duration of the viewed image, allowing us to examine how the emotional context of the current and preceding trials affected temporal judgements. We hypothesized that emotional arousal would recalibrate the weighting of prior experience during time estimation. High arousal on the current trial was expected to draw observers more strongly toward prior durations, thereby strengthening serial dependence, while low arousal should encourage more accurate encoding of the present interval. We further predicted that this bias would increase when high-arousal states persisted across consecutive trials, producing a state-dependent enhancement of temporal integration.

## Method

### Participants

Twenty-one volunteers participated in the study (13 females; mean age = 23.00 ± 1.80 years). Our goal was to examine how arousal modulates duration-based serial dependence. A recent action-manipulation study (Cheng, Chen, Glasauer, et al., 2024) reported a large effect size for the relevant slope difference (f^2^ = 0.48), and a G*Power analysis (Faul et al., 2007) indicated that at least 19 participants were needed to achieve 80% power at α = .05. All participants gave written informed consent and received compensation of €9/hour. The study was approved by the ethics committees of the Psychology Department at LMU Munich.

### Stimuli and procedure

We curated 260 images from the IAPS based on normative arousal and valence ratings. Twenty (10 neutral, 5 negative, and 5 positive) were used for practice, and 240 were reserved for the main experiment: 80 high-arousal negative, 80 high-arousal positive, and 80 low-arousal neutral (see *Supplementary 1*). Normative ratings were as follows: negative (arousal=6.84, valence=1.84), neutral (arousal=2.80, valence=5.03), and positive (arousal=5.97, valence=7.06), all *p*s < .001. Images were resized to 10.85 × 8.06° of visual angle and luminance-equated to 58 cd/m^2^, and paired with a luminance-matched mosaic image for the reproduction phase. Stimuli were presented using PsychoPy3 (Peirce et al., 2019) on a 21-inch CRT monitor (85 Hz) viewed from 60 cm in a dim, sound-attenuated room.

The study employed a duration reproduction paradigm (Figure 1A). Each trial began with a fixation (visual angle: 0.5°) for 0.5 s, followed by an image that marked the beginning of the encoding phase. The image remained visible for one of five durations (0.8, 1.0, 1.2, 1.4, or 1.6 s), which participants were instructed to memorize. After a 0.5-s blank screen, a central black dot (0.5°) signaled the reproduction phase. Participants pressed and held the down-arrow key to reproduce the encoded duration at their own pace; upon keypress, the mosaic image appeared and remained on-screen until the key was released. Participants received trial-by-trial feedback. One of five dark gray circles, arranged from left to right, appeared for 0.5 secs to indicate relative accuracy: large underestimation (< -30%), moderate underestimation ([-30%, -5%]), near-perfect reproduction ([-5%, 5%]), moderate overestimation ([5%, 30%]), and large overestimation (> 30%).

**Figure 1.**
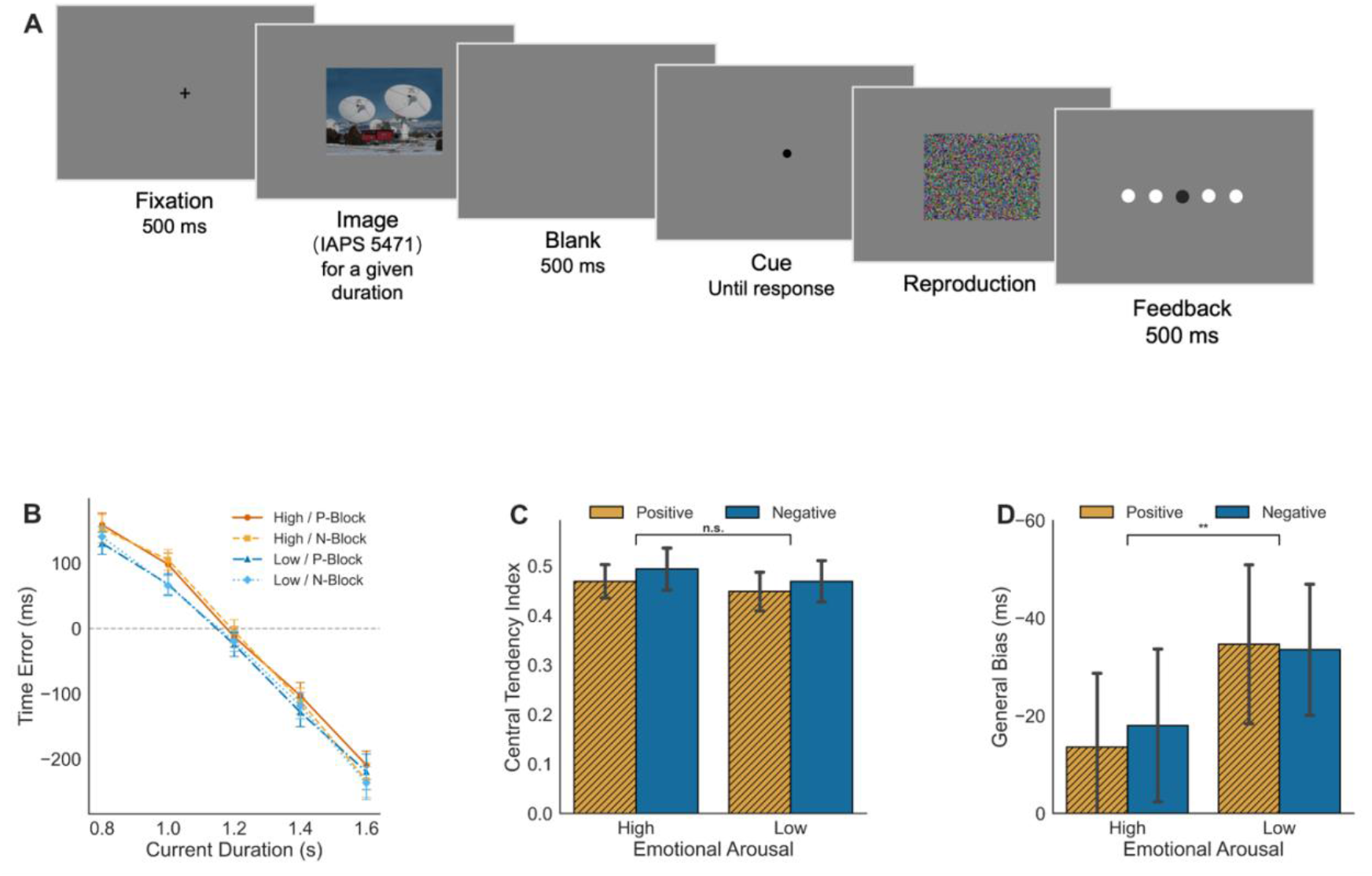
**(A)** Experimental procedure. An image (here IAPS 5471) appeared for a given duration, and participants were asked to recall the duration and then reproduce it. After reproduction, participants receive visual feedback indicating their accuracy in reproduction. (**B**) Temporal reproduction. The mean reproduction error was plotted separately for emotional valence (positive or negative) and emotional arousal (high or low). (P-Block: Positive valence; N-Block: Negative valence.) (**C**) Central tendency effect. The mean central tendency indices are plotted separately for emotional valence (positive or negative) and emotional arousal (high or low). (**D**) General biases. Error bars represent ± SEM. \*\**p* < .01.

After 20 practice trials, participants completed 640 formal trials divided into 16 blocks of 40. Blocks were split across two sessions of eight blocks each, separated by a 10-minute break. Each session contained four positive-type blocks (high-arousal positive + neutral images) and four negative-type blocks (high-arousal negative + neutral images). Eighty high-arousal images were randomly paired with 80 neutral images with each session. The same set of neutral images appeared in both block types but did not repeat within a block. To assess the influence of emotional arousal on serial dependence, we categorized trials based on the emotional arousal of the current and previous images, resulting in four transition types: High → High (HH), High → Low (HL), Low → Low (LL), and Low → High (LH). Trial sequences were balanced to ensure each transition type appeared equally often. Across all conditions, the five target durations were randomly distributed within each condition.

### Data analysis

Reproduction error was defined as the difference between reproduced and actual durations. Reaction time (RT) was measured from the onset of the reproduction cue to the initiation of the keypress. We excluded the first trial of each block and removed outliers exceeding ±3 SD from each participant’s mean reproduction error or RT, eliminating 2.11% of trials. Remaining trials were grouped by emotional block type (positive vs. negative) and arousal level (high vs. low).

#### Central tendency effect

We quantified central tendency by regressing reproduction error (*E*_n_) on the centered current duration (*Tc*_*n*_*=T*_*n*_*-1*.*2*):

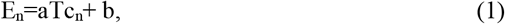

Here, the absolute slope |*a*| reflects the central tendency effect (0 = none; 1 = perfect bias), and the intercept *b* captures general reproduction bias.

#### Sequential dependence effect

To avoid confounding from the central tendency effect (see Glasauer & Shi, 2022), we measured serial dependence by regressing the current reproduction error on the previous duration after centering it— rather than using the difference between current and previous durations:

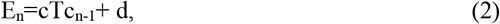

Positive slope *c* reflects attraction toward the previous duration; negative *c* indicates repulsion. The intercept *d* again represents general bias, similar to the *b* in Eq. (1).

To track how sequential effects unfolded across time, we extended the regression model to include durations from trials n-2 and n-3, and as a sanity check, from n+1 and n+2. These slopes served as indices of serial dependence. We applied Bonferroni corrections to control for multiple comparisons.

## Results

### The Central-tendency Effect

Figure 1B shows the characteristic central tendency pattern: participants overestimated short durations and underestimated long ones across all emotional contexts. The magnitude of this bias changed little with arousal or valence.

In positive valence blocks, central tendency indices averaged 0.469 ± 0.034 for high-arousal trials and 0.448 ± 0.039 for low-arousal trials. In negative valence blocks, the corresponding values were 0.494 ± 0.043 and 0.467 ± 0.042 (Figure 1C). Each index differed reliably from zero (*t*s > 11.03, *p*s < .001), confirming a consistent central tendency effect. A two-way repeated-measures ANOVA revealed no significant main effects of Valence and Arousal, nor their interaction (all *F*s < 2.274, *p*s > .147, p2 < .102). This consistency likely stems from the identical duration range used in all conditions, which may have fostered a stable temporal prior.

We then assessed the general reproduction bias (*b*). Participants tended to underestimate durations in all contexts, but high arousal attenuated this bias. In positive blocks, underestimation averaged 14 ± 15 ms for high-arousal stimuli and 35 ± 16 ms for low-arousal stimuli; in negative blocks, the corresponding values were 18 ± 16 ms and 34 ± 13 ms (Figure 1D). A two-way ANOVA revealed a significant main effect of Stimulus Arousal (*F*_*(1,20)*_ = 13.56, *p* = .001, p2 = 0.404), with no effect of Valence and no interaction (*F*s < 0.51, *p*s > .48). Across valence conditions, high arousal shifted perceived duration upward, reducing the overall underestimation.

#### The Serial-Dependence Effect

We estimated serial dependence by regressing the current reproduction error on the duration of the preceding trial. A three-way ANOVA on the resulting slopes showed no valence effects (see Supplementary 2); therefore, the analyses below are collapsed across valence.

Figure 2A illustrates a clear sequential attraction: reproduction errors increased with the previous duration in every arousal combination. All slopes were significantly greater than 0 (*t*s > 2.787, *p*s < 0.011), confirming robust attraction toward prior durations.

**Figure 2.**
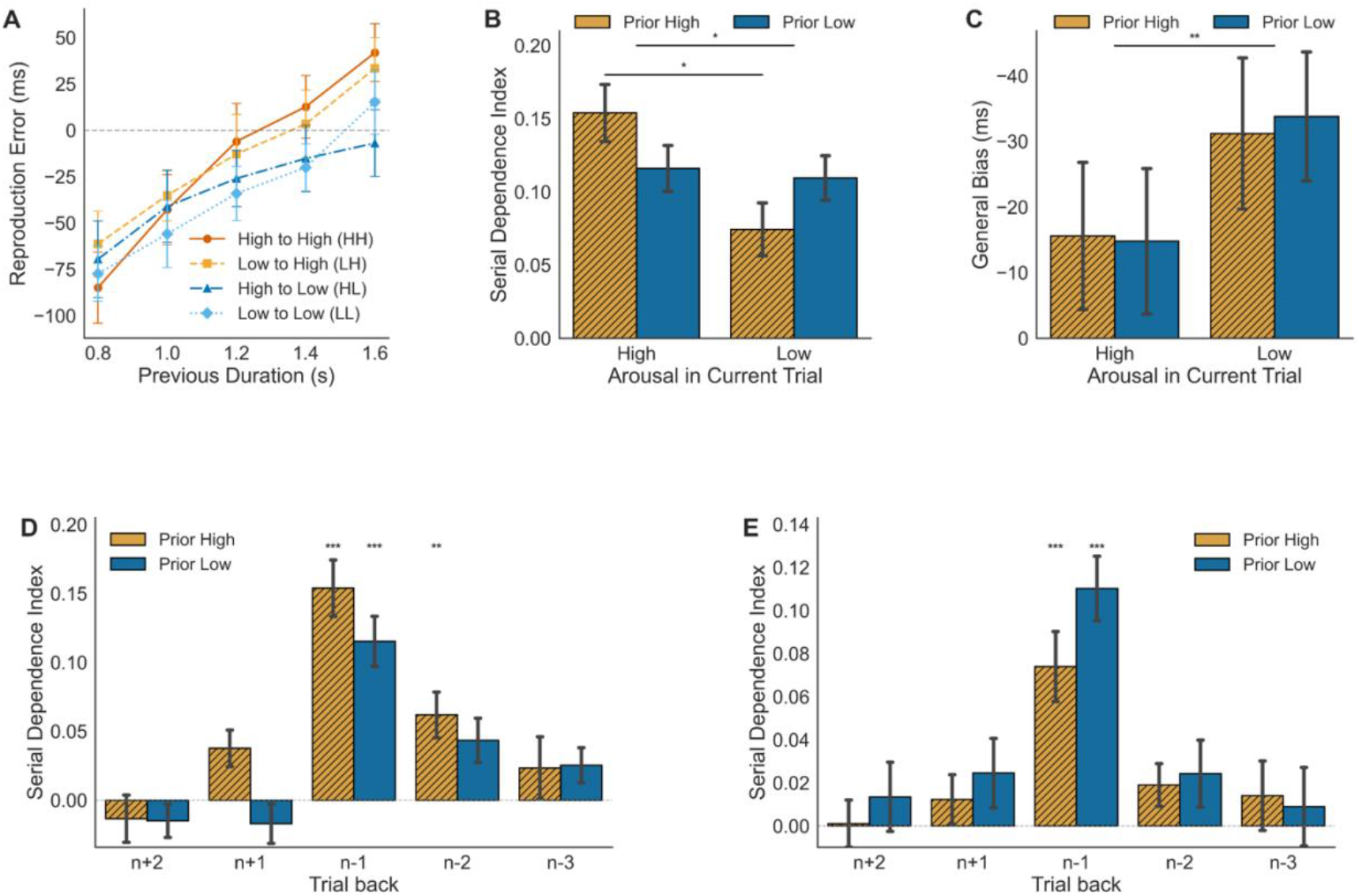
Serial dependence effect. (**A**) Mean reproduction errors are plotted as a function of previous durations for the HH, LH, HL, and LL conditions. (**B**) The serial dependence indices are shown separately for current high and low arousal conditions, with comparisons based on whether the previous trial’s arousal was high or low. (**C**) General biases. (**D**) Serial dependence indices for the current high arousal trials, plotted separately for the prior high arousal and low arousal conditions in the preceding trials (n-1, n-2, and n-3) and in the future trials (n+1, n+2). (**E**) Serial dependence indices for the current low-arousal trials. Error bars represent ± SEM. \*\*\**p* < .001, \*\**p* < .01, \**p* < .05.

A two-way ANOVA on slopes revealed that Current Arousal significantly strengthened serial dependence (*F*_*(1,20)*_ = 7.356, *p* = .013, p2 = 0.269), whereas Previous Arousal alone did not (*F*_*(1,20)*_ = 0.01, *p* = .92, p2 = 0.001). More intriguingly, the Current × Previous Arousal interaction was significant (*F*_*(1,20)*_ = 7.492, *p* = .013, p2 = 0.273), suggesting that serial dependence is not simply a carryover from the past - it intensifies when both current and prior stimuli are emotionally arousing (see Figure 2B). Post-hoc comparisons confirmed this: when both current and previous trials were high-arousal, serial bias peaked (*t*_*(20)*_ = 2.450, *p* =.047, *BF*_*10*_ = 2.485, Bonferroni-corrected). General biases across these conditions mirrored the pattern observed in Figure 1D (Figure 2C).

We next examined how far this arousal-driven bias extended. Regressions incorporating durations from trials n-1, n-2, and n-3, and for verification, from future trials n+1 and n+2 (Figure 2D and E). In current high-arousal conditions, the serial dependence extended beyond the immediately prior trial: when the prior two trials (n-2) were also high in arousal, a significant influence was still detectable (slope: 0.062 ± 0.017, *t*_*(20)*_ = 3.723, *p* = .007, Bonferroni-corrected). This extended effect did not appear in low-arousal contexts, suggesting a specific amplifying role for sustained emotional intensity. No evidence of bias from future trials emerged (*ps* > .067).

## Discussion

This study examined the impact of emotional states on serial dependence in time perception. By inducing high-arousal (positive or negative) and low-arousal (neutral) states, we found that emotional arousal not only lengthened perceived durations but also strengthened sequential biases. Serial dependence grew especially pronounced when high-arousal trials followed other high-arousal trials, whereas emotional valence exerted no measurable influence. These findings position arousal as a key internal factor that governs how the perceptual system balances present inputs with recent experience.

The robust lengthening of perceived duration during high arousal aligns with internal clock models (Droit-Volet & Gil, 2009; Shi et al., 2013), which propose that arousal accelerates pacemaker rate and increases the number of temporal “pulses” accumulated for a given interval. When compared against a reference memory, the inflated pulse count makes the interval feel longer.

What’s new here is the link between arousal and serial dependence, showing arousal also modulates the integration of past and present time information. One possible explanation for stronger serial dependence under current high arousal is attentional narrowing. Elevated arousal is associated with intensified focus on emotionally salient content (Lam et al., 2020), which might reduce encoding precision for the current duration and encourage heavier reliance on the recent past. However, several features of our data challenge a pure distraction account. First, if high-arousal images had caused distraction, which should shorten perceived duration, yet high-arousal intervals were consistently overestimated. Second, degraded encoding precision should increase central tendency, but central-tendency slopes remained stable across arousal conditions. Third, a generic attentional lapse cannot account for the sharp amplification of serial dependence specifically in High-High transitions. If distraction were the primary driver, high-arousal trials should show reduced precision regardless of the preceding trial. Instead, sequential biases intensified only when elevated arousal persisted across trials. These patterns indicate that arousal does not simply impair temporal encoding—it alters how the system integrates recent experience into current judgments.

A broader theoretical lens situates these results within adaptive models of perception that treat serial dependence as a stabilizing mechanism under uncertainty (Fischer & Whitney, 2014). Moment-to-moment fluctuations in arousal may act as an internal source of noise, prompting the system to rely more heavily on past information when current inputs become less dependable. This interpretation aligns with recent work showing that external stimulus uncertainty–such as reduced contrast or lower reliability–boosts serial dependence (Ceylan et al., 2021). Our findings suggest that emotional arousal produces a similar functional consequence: when internal states heighten volatility or energetic load, the perceptual system compensates by granting recent experience greater weight. The extended influence observed up to two trials back under sustained high arousal further supports the idea that internal arousal states magnifies the temporal window over which past information is integrated.

Importantly, our results converge with emerging evidence that serial dependence is driven by the quality of the *current* sensory or cognitive state rather than the previous one. Gallagher and Benton (2022) and Ozkirli and Pascucci (2023) each showed that serial dependence in orientation judgments increases when the *current* stimulus is uncertain, whereas uncertainty in the previous stimulus has little impact. Ozkirli and Pascucci (2023) argued that these effects reflect a shift in the observer’s internal processing regime: serial dependence increases when the system enters a state characterized by reduced confidence, heightened volatility, or elevated internal noise. We find a strikingly parallel pattern: high arousal in the current trial strengthened sequential attraction, but previous-trial arousal alone did not. Even more compelling, both domains show the same continuity signature—serial dependence peaks when high-intensity states persist from one trial to the next (high → high). This continuity signature suggests a shared processing rule across perceptual domains: internal states set the operating mode of the perceptual system, and this mode determines how strongly recent history shapes the present.

This interpretation also departs from simple Bayesian expectations. A Bayesian observer should downweight unreliable past inputs and rely more on the current stimulus when the past becomes noisy (Fritsche et al., 2020). Instead, we observed the reverse pattern: when arousal was high—an internal state that could plausibly degrade the precision of both current and previous percepts—the influence of the past intensified. This does not necessarily violate Bayesian principles, but it highlights how the observer’s internal state dynamically reshapes the effective prior or the integration strategy. From this perspective, arousal shifts the system into a processing mode that privileges temporal continuity, granting recent experience more authority in guiding the present. This framework invites future models to incorporate internal states, such as arousal, as gating mechanisms that regulate how strongly past information constrains ongoing perception.

Together, these findings support a state-dependent account of serial dependence. The weight assigned to past information is not fixed but is continuously recalibrated by internal states such as arousal, which shape the perceptual system’s operating regime. In high-arousal contexts—where perceptual, mnemonic, and attentional resources may be strained or hyperactivated—recent stimuli gain salience and exert a stronger pull on current judgments. This state-dependent modulation reveals a form of perceptual inertia: continuity in internal state fosters continuity in perception.

In sum, emotional arousal both distorts the perception of individual intervals and reshapes how past experience guides present judgments. The sequential biases emerged when high-arousal states persisted across trials, demonstrating how internal states govern the bridge between past and present in time perception.

## Data Availability

All data needed to reproduce our study analyses are available online in the following repository: https://github.com/msenselab/emotion-and-serial-dependence

## Supplementary Materials

**Arousal-Driven Serial Dependence: Internal States Modulate Perceived Duration**

**Supplementary 1: IAPS Picture Numbers Used in the Experiment**

This supplementary material provides the complete list of International Affective Picture System (IAPS; Lang, Bradley, & Cuthbert, 2008) picture numbers used in the formal experiment. These stimulus identifiers enable exact replication of the experimental materials.

Stimulus Selection Criteria

Pictures were selected from the IAPS database based on normative ratings of arousal and valence. We selected images that fell into three distinct emotional categories to manipulate arousal and valence orthogonally:

- **Neutral pictures**: Low arousal, neutral valence
- **Positive pictures**: High arousal, positive valence
- **Negative pictures**: High arousal, negative valence

Each category contained 80 unique images, totaling 240 IAPS pictures used in the experiment.

1. **Neural IAPS Picture Numbers (N=80)** 7175, 7004, 7010, 7000, 7020, 7490, 7491, 7950, 7031, 7060, 7187, 2190, 2381, 7059, 7041, 2480, 2840, 7217, 2397, 7150, 7052, 7006, 7110, 5130, 7080, 7185, 7100, 7035, 9360, 5471, 5740, 7705, 2440, 2038, 2580, 5510, 5530, 4279, 6150, 7179, 2890, 7040, 7050, 7053, 2102, 7090, 7235, 2393, 5520, 7034, 7056, 7233, 2570, 7025, 7205, 2499, 7224, 7700, 5731, 2221, 2305, 7234, 2980, 7160, 7500, 2506, 7038, 2870, 7547, 8311, 7043, 7055, 7161, 7140, 2880, 5533, 5500, 7036, 7130, 2513
2. **Positive IAPS Picture Numbers (N=80)** 9156, 8502, 8501, 8500, 8499, 8496, 8490, 8470, 8460, 8400, 8380, 8370, 8300, 8210, 8200, 8193, 8190, 8186, 8185, 8180, 8179, 8178, 8170, 8161, 8130, 8116, 8090, 8080, 8041, 8040, 8033, 8031, 8021, 7502, 7330, 7270, 7220, 5910, 5626, 5623, 5470, 5460, 5450, 5270, 4694, 4690, 4677, 4676, 4658, 4653, 4650, 4645, 4643, 4640, 4609, 4607, 4599, 4598, 4572, 4561, 4550, 4542, 4538, 4537, 4535, 4533, 4532, 4531, 4520, 4510, 4503, 4490, 4470, 4460, 2389, 2209, 2208, 1722, 1650, 4689
3. **Negative IAPS Picture Numbers (N=80)** 3080, 6230, 3170, 9410, 6350, 3053, 3120, 3010, 3266, 3130, 3010, 3060, 3069, 3064, 2811, 6313, 6550, 3063, 3500, 6510, 3102, 6540, 3030, 3400, 3071, 3068, 3100, 3110, 3140, 3150, 2730, 9252, 1052, 1201, 9921, 1525, 6560, 6300, 6312, 6360, 3530, 5971, 9405, 6260, 1932, 6210, 6250, 9433, 6830, 9600, 6315, 9250, 6834, 9050, 6200, 6821, 6370, 6212, 6415, 9810, 9571, 9570, 9040, 6570, 2683, 9042, 9910, 6530, 9622, 6243, 9620, 2981, 9254, 9400, 3225, 6021, 6022, 8485, 9911, 3168

**Supplementary 2: Three-way repeated-measures ANOVA on serial dependence**

We estimated serial dependence by regressing the current reproduction error on the duration of the previous trial within each Valence, Current Arousal, and Previous Arousal condition. The resulting slopes served as indices of serial dependence. A three-way ANOVA on these slopes revealed no significant interaction among Valence, Current Arousal, and Previous Arousal (F_(1,160)_ = 0.292, p = .590). When we focused on arousal-related effects, a clearer pattern emerged. There was no main effect of Valence (F_(1,160)_ = 0.030, p = .863) and no effect of Previous Arousal (F_(1,160)_ = 0.007, p = .936), but Current Arousal exerted a significant influence (F_(1,160)_ = 6.296, p = .013, η^2^_p_ = 0.038): participants showed stronger serial dependence when the current stimulus elicited high arousal.

A Current × Previous Arousal interaction also emerged (F_(1,160)_ = 4.527, p = .035, η^2^_p_ = 0.028), indicating that serial dependence does not simply carry over from the preceding trial. Instead, it intensifies when both the current and prior stimuli are emotionally arousing. Post hoc comparisons supported this pattern: serial bias peaked when both current and previous trials were high in arousal (t_(20)_ = 2.450, p = .047, BF_10_ = 2.485, Bonferroni-corrected).

